# Olive mill wastewater extract as a potential mosquito larvicide

**DOI:** 10.1101/2022.01.18.476713

**Authors:** Maram Halabi, Alon Silberbush, Hassan Azaizeh, Ben Shahar, Eyal Kurzbaum

**Affiliations:** Department of Biology and Environment, Faculty of Natural Sciences, University of Haifa – Oranim, Israel; Institute of Applied Research, The Galilee Society, P.O. Box 437, Shefa-Amr 20200, Israel; Tel Hai College, Department of Environmental Science, Upper Galilee 12208, Israel; Department of Natural Resources & Environmental Management, University of Haifa, Haifa, 3498838, Israel; Shamir Research Institute, University of Haifa, P.O.B. 97, Qatzrin, Israel; Department of Geography and Environmental Studies, University of Haifa, Mount Carmel, Haifa 3498838, Israel

**Keywords:** Olive mill waste water, *Culex laticinctus* (Edwards) larvae, larvicide, Lethal concentrations, Sub-lethal concentrations.

## Abstract

The larvicidal potential of olive mill waste water (OMWW) extract against mosquito larvae was evaluated. We exposed 2^nd^ and 4^th^ instar *Culex laticinctus* (Edwards) larvae to increasing OMWW concentrations. In addition, the effects of sublethal OMWW concentrations on larval development time and adult size were tested as well. The larvicidal activity of OMWW extract showed LC_50_ values of 10.08 and 50.07 parts per thousand (ppt) against the 2^nd^ and 4^th^ instars respectively. Larvae that developed in 1 ppt OMWW solution showed sex-specific responses. Males prolonged time to pupation, while females exhibited size reduction in comparison to controls. These results show that OMWW, which is known as agricultural byproduct waste, may be reused as a biopesticide.

## Introduction

The olive oil industry is one of the most important branches of agriculture in the Mediterranean basin producing over 90% of the global olive oil production (IOC 2022). This industry also generates solid and liquid waste byproducts characterized by their dark color and a typical odor, which are not easily degradable. Solid waste can be recycled and used as an ingredient of several products such as fertilizers, ethanol and high value-added biomolecules (Muscolo et al. 2021) or lightweight aggregates (Moreno-Maroto et al. 2019). On the other hand, the liquid byproduct, olive mill wastewater (OMWW), is significantly more difficult to treat. This liquid is characterized by high concentrations of polyphenols and tannins, in addition to low pH and high chemical and biochemical oxygen demands. These qualities inhibit biological decomposition and place OMWW as one of the most contaminating effluents among those produced by the agrofood industries (Messineo et al. 2020).

Previous studies suggested that the high concentrations of phenols, acids and sugar derivatives associated with OMWW may be applied to be used as biofertilizers, environmentally crop protections, ethanol and high value-added biomolecules (Azzaz et al. 2020). These qualities of OMWW also make its extractions to be potentially useful biopesticides. Some studies have shown that extracts of OMWW have the potential to inhibit the growth of bacteria, fungi and weeds (El-Abbassi et al. 2017). However, only limited work was performed exploring the potential of OMWW extracts to inhibit the growth or kill insects and other arthropods. The existing data on the insecticidal potential of OMWW focuses on phytophagous insects (Larif et al. 2013, Lahcene et al. 2018, Boutaj et al. 2020).

The main objective of the current study was to examine the biopesticide potential of OMWW extracts against the development and survival of *Culex* larvae.

## Methods

### Larval Collection

*Culex laticinctus* (Edwards) egg rafts were collected from plastic ovitraps placed at Oranim College Campus, Tivon, Israel. Egg rafts collected from the water were placed in sperate plastic cups. Sampled larvae from each batch were reared to 4^th^ instar and identified to species (Becker et al. 2010). First instar larvae were transferred into 500 ml plastic cups containing aged tap water.

### OMWW extract

OMWW for this study was obtained from a nearby olive mill press (Iksal, Galilee region, Israel). The OMWW was treated with 20% ethanol (v/v) and stored at 4°C until use within few weeks. One liter was centrifuged (8000g for 10 min), supernatant was then filtered through a filter paper and evaporated under vacuum using the Rotary evaporator, until it reaches a volume of 250 ml. 250 ml of 95% ethanol solution was added to obtain two steps: solid precipitate (cellulose) and liquid layer (dissolved polyphenols). The solid precipitate was removed from the mixture by filtration, and the liquid fraction was evaporated at 40 °C, using Rotary evaporator to a final volume of 250 ml. The process was repeated until no more of solid fraction was obtained from the mixture. Subsequently, the resulting liquid was evaporated under vacuum to form a 10 mg substance (resembling a dark ointment) called an anti-solvent solution. The extract was kept at 4 °C. The anti-solvent solution was mixed with distilled water and stirred for 16 hours at room temperature. The solution was then centrifuged, and the filtered solution was adjusted to a final concentration of 145 ppt (parts per thousand).

### Larval toxicity bioassays

Toxicity tests were performed by using a standard bioassay (World Health Organization 2005) on 2^nd^ and 4^th^ instar larvae to determine LC_50_ value. Larval siblings from the same batch were placed in cups containing increasing concentrations of OMWW extract. Second instar larvae were introduced into 100 ml of aged tap water (25 per cup) with the concentrations of 5, 10, 15, 20 and 25 ppt OMWW in addition to control cups without OMWW. Each concentration was repeated 3 times. For the larger and more resistant 4^th^ instars we used cups with 40 ml of water containing 3 larvae. These larvae were exposed to concentrations of 2, 20 and 50 ppt OMWW in addition to control. Control treatment was repeated 3 times, 2 ppt 10 times, 20 ppt 3 times and 50 ppt, 6 times. Larvae were fed with a finely grounded mixture of fish flakes (42.2% crude protein, Sera-Vipan, Heinsberg, Germany) and rodent chow (17% protein, Ribos, Haifa, Israel). Cups were kept at a temperature of 24.83 ± 0.84 °C (mean ± SD). We recorded larval mortality after 48 hours of incubation.

### Sublethal effects of OMWW on larvae

Twenty sibling larvae were transferred into 200 ml plastic cups containing 100 ml aged tap water within 24 hours of hatching. Each block consisted of a cup containing 1 ppt OMWW solution (equivalent to 10% 2^nd^ instar LC_50_) and aged water control (4 blocks in total). Larvae were fed every three days with 0.05 ± 0.003 (mean ± SD) grams of the mixture used in the previous experiment. Cups were kept at a temperature of 24.83 ± 0.84 °C (mean ± SD). Pupae were kept in separate test tubes until emergence. Emerging adults were placed in a drying oven at 50 °C, for 24 hours and were identified to sex. We separated the right wing of each individual. Wings were photographed and length measured from axillary incision to the apex (figure 1) using Dino-lite (DinoCapture 2.0). We recorded the number of days to pupation, nuber of emerging adults and the average wing length.

**Figure 1:**
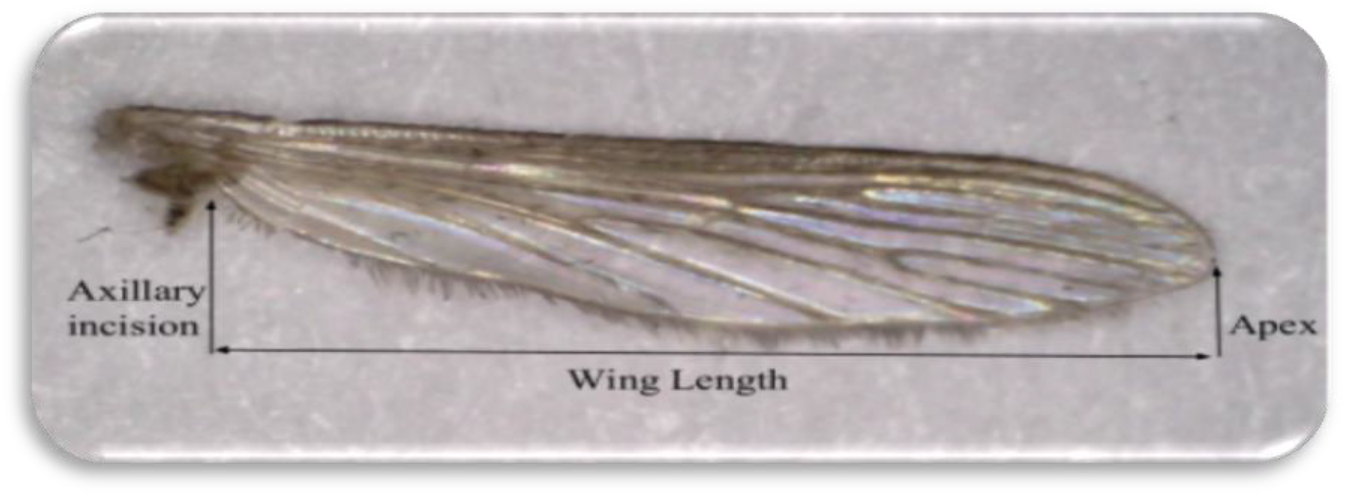
Length measurement of *Culex laticinctus* wing

## Statistical analysis

LC_50_ values at 95% confidence intervals values were calculated using Probit analysis (Finney 1952). We used Pearson goodness of fit test in order to evaluate observed distribution. A heterogeneity factor was used to calculate confidence limits when model assumptions were not met (p<0.15). The effects of sublethal concentrations on larval time to pupation and adult wing length were analyzed separately using Linear Mixed Models (LMM), with the variables “Sex” and “Treatment” (OMWW and control) as fix factors and “Block” as a random factor. These analyses considered random sampling (Block effect), and the fact that male mosquitoes usually pupate faster and are smaller in size than females (Sex effect). All analyses used SPSS statistics for windows version 24 with Type III sums of squares (IBM 2016).

## Results

Second instar larvae showed 100% mortality after being exposed to 25 ppt OMWW for 48 hours. Fourth instar larvae did not exhibit 100% mortality when exposed to the maximal tested concentration of 50 ppt OMWW. Probit analysis found the LC_50_ value for 2^nd^ instar to be ~5 times lower than that of 4^th^ instars. Slope ratio showed mortality rates for 2^nd^ instars to be 3.25 times higher in comparison to 4^th^ instars (Table 1).

**Table 1:**
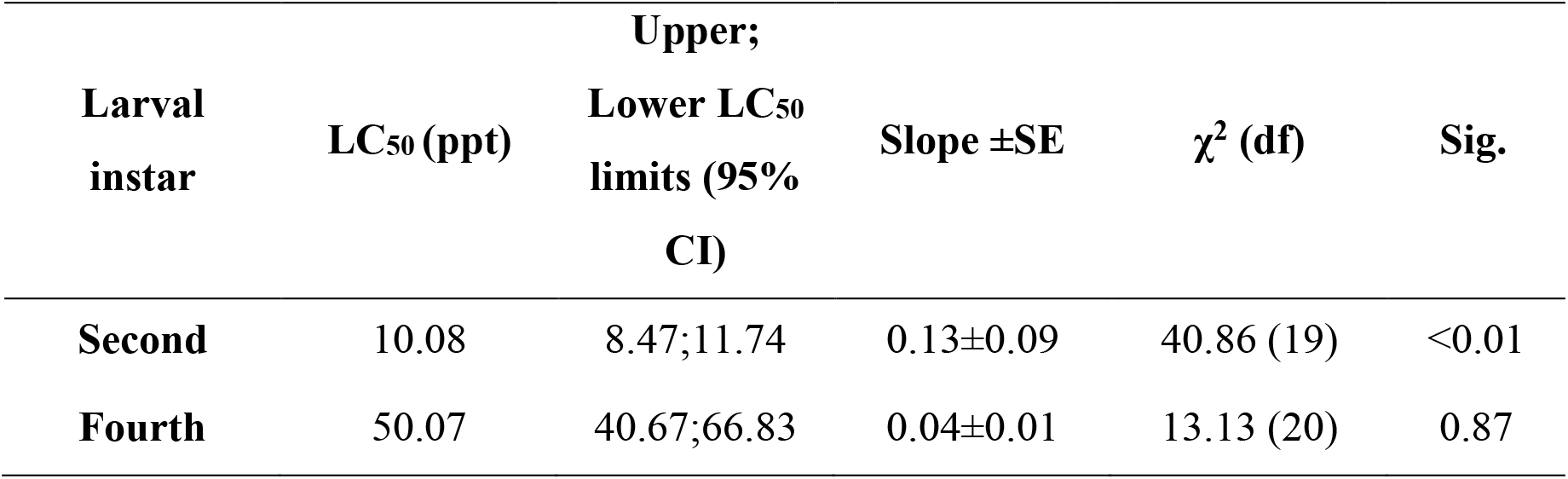
Probit analysis results for LC_50_ values, slopes of the linear models and heterogeneity test results for 2^nd^ and 4^th^ instar *C. laticinctus* larvae exposed to increasing OMWW extract concentrations for 48 hours.

The number of adults emerging from solutions containing 1 ppt OMWW did not differ from the number of adults emerging from control cups (paired t-test: t_3_=0.52; p=0.64 and t_3_=0.42; p=0.7 for males and females, respectively). When observing the effects of the different factors on time to pupation we found a strong sex effect, i.e. males pupated faster than females (F_1,147.13_=27.9; p<0.001). Female pupation time did not vary among the treatments (Figure 2a), however males exposed to OMWW prolonged time to pupation resulted in a significant “Sex × Treatment” interaction (F_1,147.08_=6.4; p<0.001). The overall effect of OMWW by itself was not statistically significant (F_1,147.04_=1.5; p=0.23). Wings of adult females were significantly longer than those of males (Sex effect: F_1,147.13_=206.9; p<0.001). We also found that females developed in OMWW had shorter wing length than their siblings developed in control water. This trend was reversed for males whose wings were longer in the OMWW treatment resulting in a significant “Sex × Treatment” interaction (F_1,147.3_=18.9; p<0.001). The overall effect of OMWW by itself was not statistically significant (F_1,147.13_=1.04; p=0.31).

**Figure 2:**
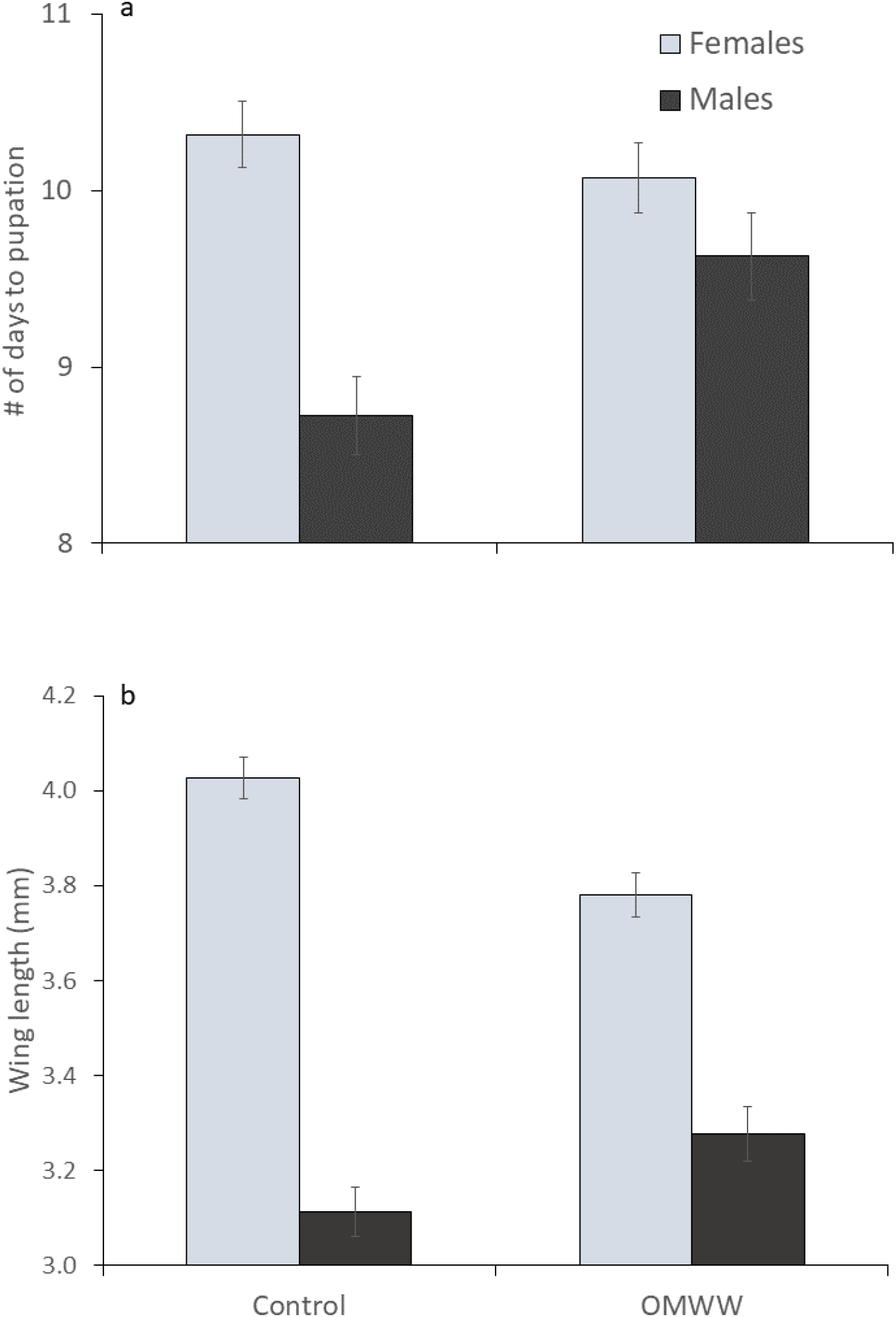
(a) Number of days to pupation for male and female larvae treated with 1 ppt OMWW solution and control, (b) Adult wing length in mm (at the bottom how it was measured) for male and female larvae treated with 1 ppt OMWW solution and control. Error bars denote ± 1 SE.

## Discussion

Results showed that OMWW extract is indeed toxic to *C. laticinctus*. Early larval instars are often more sensitive to insecticides than late instars (Mulla 1961), hence efficiency is usually measured for late instars. *Culex laticinctus* is one of the most common mosquito species in the Mediterranean region, and their breeding sites often consist of small, artificial water bodies (Becker et al. 2010). Hence, we may assume that larval *C. laticinctus* are often exposed to olive foliage and are relatively resistant to OMWW. Therefore, we expect that OMWW extracts should be more lethal for larval mosquito species that are not associated with that region.

The LC_50_ level of OMWW for 4^th^ instars was shown to be about 50 ppt (Table 1). We can expect that the lethal effect of OMWW extracts will increase further by removal of some of the compounds by different types of fractionation, a practice where bioactive compounds are separated from the crude extract (Tafesh et al. 2011). For example, fractions containing mainly polyphenol from OMWW cause mortality to *Euphyllura olivina* and *Aphis citricola* by direct spraying of liquid solutions. The LC_50_ in these studies was recorded at 0.36 and 2.12 ppt for the two Hemipterans respectively (Larif et al. 2013).

Our results also show the effects of sublethal concentrations of OMWW. Low concentrations of pesticide may reduce pest population fecundity over time without causing immediate mortality. For example, inhalation of OMWW extracts by pupae of the Mediterranean flour moth *Ephestia kuehniella* (Lepidoptera: Pyralidae) resulted in extended pupal duration period. In addition, emerging adults delayed time to oviposition and oviposited less eggs over fewer days (Lahcene et al. 2018). Crude extract of OMWW caused weight loss for larvae of the palm tree pest *Potosia opaca* (Coleoptera: Scarabeidae). Increased larval mortality was recorded well over a week following multiple treatments (Boutaj et al. 2020).

The effects of sublethal OMWW concentration was not equally visible for males and females. Males extended their time to pupation by almost a full day while females were not significantly affected. This resulted in an almost simultaneous emergence of males and females from OMWW treated cups (Figure 2a). By contrast, the same males emerging from OMWW treated cups had longer wing on average while female wings were shorter (Figure 2b). The two analyses results are likely not independently from one another. In most mosquito species males have a shorter larval stage, emerge 1-2 days before the females and the adult males are significantly smaller than the females (Becker et al. 2010). However, males become sexually mature only 1 day after emergence, at the same time as later emerging female of the same cohort (Becker et al. 2010). A situation where males and females emerge simultaneously following exposure to OMWW may result in delayed mating and a reduction of the overall the number of mating couples. For a majority of mosquito species fecundity is strongly correlated with wing length where larger females will have longer wings and produce more eggs (Vinogradova 2000). Thus, a population where females who are smaller in size following exposure to OMWW (Figure 2b), will likely produce a reduced number of offspring.

Overall, our results support the possibility that OMWW may be applied as a source for mosquito larvicide. Future research should focus on fractionation, isolation and identification of compounds from different fractions of OMWW with strong insecticidal activity. It is important to emphasis that this approach suits well the circular bio-economy and green chemistry models that deal in the importance of valorization of agro-wastes and by-products generated by agricultural and agro-industries.

## Acknowledgments

Yoram Gerchman and Nimrod Shteindel assisted in data collection. Avi Bar-Massada, and Elad Chiel helped with several aspects of this study. This work was supported by the Margolin grant awarded to Maram Halabi.

